# Three membrane fusion pore families determine the pathway to pore dilation

**DOI:** 10.1101/2022.02.23.481713

**Authors:** Rui Su, Shuyuan Wang, Zachary McDargh, Ben O’Shaughnessy

**Affiliations:** Department of Chemical Engineering, Columbia University, New York City, NY, USA; Department of Physics, Columbia University, New York City, NY, USA

**Keywords:** membrane fusion, fusion pore, membranes, exocytosis, soap films, catenoid

## Abstract

During exocytosis secretory vesicles fuse with a target membrane and release neurotransmitters, hormones or other bioactive molecules through a membrane fusion pore. The initially small pore may subsequently dilate for full contents release, as commonly observed in amperometric traces. The size, shape and evolution of the pore is critical to the course of contents release, but exact fusion pore solutions accounting for membrane tension and bending energy constraints have not been available. Here we obtained exact solutions for fusion pores between two membranes. We find three families: a narrow pore, a wide pore and an intermediate tether-like pore. For high tensions these are close to the catenoidal and tether solutions recently reported for freely hinged membrane boundaries. We suggest membrane fusion initially generates a stable narrow pore, and the dilation pathway is a transition to the stable wide pore family. The unstable intermediate pore is the transition state that sets the energy barrier for this dilation pathway. Pore dilation is mechanosensitive, as the energy barrier is lowered by increased membrane tension. Finally, we study fusion pores in nanodiscs, powerful systems for the study of individual pores. We show that nanodiscs stabilize fusion pores by locking them into the narrow pore family.

**Significance:** During neurotransmission, hormone release and other fundamental processes, secretory vesicles fuse their membranes with target membranes to release contents through an initially small membrane fusion pore that subsequently dilates. Dilation is assisted by proteins such as SNAREs and synaptotagmin. While macroscopic soap film shapes are well characterized, finding exact solutions for microscopic cellular membrane surfaces is made more complex by bending energy constraints. Here, computational analysis revealed three families of fusion pores between two membranes. Our work suggests membrane fusion generates a member of the narrow pore family, and pore dilation is a transition to the wide pore family. The energy barrier that SNAREs or synaptotagmin must surmount to achieve dilation is set by a third unstable intermediate pore family.

## Introduction

Phospholipid membranes are essential to life as enforcers of compartmentalization, providing impermeable yet flexible surfaces whose highly adaptable shapes readily adjust to enclose compartments and maintain specialized conditions and contents within (1–3). Consisting of ~5nm thick phospholipid bilayers dressed with transmembrane proteins, channels, receptors and other machineries (4, 5), cell membranes are used to enclose diverse compartments, from ~ 50 nm synaptic vesicles for neurotransmitter release to ~ 100-400 nm dense core vesicles for secretion of hormones such as insulin (6–8), to large organelles (9) and the cell itself.

The most extreme demands on membrane shape occur during processes such as secretion and trafficking, when membranes of different compartments undergo topological change by membrane fusion, resulting in a membrane fusion pore connecting the compartments (10). Here we consider purely lipidic fusion pores (11), but the initial fusion pore in cells has also been proposed to be proteinaceous, lined by a complex of transmembrane domains of the SNARE proteins syntaxin and synaptobrevin (12–15). The simplest membrane shape is the sphere, a surface of minimal area that also minimizes the membrane bending energy, adopted by closed secretory vesicles. Much less is established about the shapes and dynamics of fusion pores.

At neuronal synapses, the cellular fusion machinery creates a fusion pore within less than a millisec of presynaptic calcium influx, inferred from delay times in electrophysiological measurements of excitatory postsynaptic currents (16). Conductance measurements at the Calyx of Held revealed minimum synaptic vesicle pore diameters of ~ 1 nm that can flicker repeatedly. A historic development was the use of microelectrodes to detect the far slower release from non-synaptic secretory cells which showed that the initial fusion pore is very small but often dilates dramatically after a delay (17). In amperometry recordings this is indicated by a flickering low amplitude 1-50 ms “foot” signal that abruptly transits to a spike (18–20). In neuroendocrine chromaffin cells, pores can dilate to ~ 100 nm and remain open for >30 sec (21, 22). Fusion pores are mechanosensitive, as higher membrane tension correlates with larger pores and favors full fusion and contents release over partial release (“kiss-and-run”) (23, 24).

The sizes, shapes and evolution of fusion pores are critical to exocytosis, when secretory vesicles fuse with target membranes and release contents through pores. The narrow initial pore may serve as a molecular sieve to modulate size and rate of vesicle cargo release (25, 26), while full release requires the pore to dilate. For example, fusion events at synaptic terminals may result in partial or total neurotransmitter release, depending on whether the nascent pore dilates fully or reseals, with important consequences for synaptic activity (27, 28). Pore dilation is assisted by SNARE proteins and Synaptotogamin (Syt) (29–31), components of the cellular fusion machinery (32–34), but the membrane energy barriers they must overcome to dilate pores are not known. Impaired pore expansion is associated with disease such as type 2 diabetes where insulin release is misregulated (35). Membrane fusion pores are also used for host cell entry by membrane-enveloped viruses such as influenza, HIV and SARS coronaviruses, whose genomes enter via a fusion pore following fusion of the viral and host membranes by specialized glycoproteins (36–38).

Theoretical works have sought to establish fusion pore shapes between two planar membranes, accounting for free energy contributions from membrane tension (tending to minimize area) and from surface bending energy (39). Fusion pores were predicted, but the shapes were assumed to belong to a particular class, either toroidal, ellipsoidal, or a combination of catenoidal and cylindrical (40–42), or were approximated by polynomials (41, 43). Continuum models and MARTINI simulations showed pores that bowed outwards beyond the planar membranes (43, 44). These studies identified a single family of narrow fusion pores, with minimum diameter < 10 nm. By contrast, an important recent study found exact solutions minimizing the Helfrich free energy corresponding to three fusion pore families, using freely hinged boundary conditions at the membrane edges (45). Membrane tethers are structures closely related to fusion pores, typically pulled from cells or artificial liposomes by optical tweezers that use the measured pulling force to infer membrane tension (46, 47). Rigorous analysis showed that long tethers approach a cylindrical shape, with a more complex transition region where the tether attaches to the planar membrane (48, 49).

Classical soap films are a natural reference point for the study of microscopic membrane surfaces. These macroscopic surfactant-stabilized water films have surface tension and thus if equilibrated adopt surfaces minimizing the area, including the spherical soap bubble. Another classical shape is the catenoid. Given two planar soap films enclosed by hoops, if the films become fused the preferred shape is the catenoid, the minimal area open surface given the constraints (50). In fact two catenoidal solutions exist, but only the wider catenoid family is stable. Bending energy is unimportant for macroscopic soap films, but for the microscopic scales of cell membranes this component becomes significant so the selected surface shape minimizes the sum of the tension and bending energies, the Helfrich free energy (39). A natural question is whether microscopic membrane fusion pores adopt catenoid or catenoid-like solutions similar to soap films. The study of ref. (45) showed that when bending energy features, the second thinner catenoid family can also be realized if the far field boundary conditions are freely hinged (45).

Here we obtain exact shapes and energetics of fusion pores between planar membranes by minimizing the Helfrich free energy (39). With simplified freely hinged boundary conditions, appropriate to soap films, there are three families as reported by Powers et al. (45): the wide and thin catenoids and the tether-like family. We show the thin catenoid is unobservable in soap films, being locally stable for membrane tensions within a band whose width decreases with increasing membrane size. Accounting properly for bending constraints from the planar membranes, we find three fusion pore families with shapes and energetics close to those of the freely hinged families. We propose that the nascent fusion pore generated by the cellular fusion machinery during exocytosis is the quasi-thin catenoid, whose dilation is a transition to the quasi-wide catenoid as observed in amperometric foot-to-spike transitions (18–20). The transition energy barrier is set by the unstable tether intermediate and is lowered by membrane tension, consistent with experiments showing facilitated dilation at higher tensions (23, 24). Finally, we study nanodiscs, model systems allowing long time measurements of stabilized pores (30, 31). We show that nanodiscs stabilize fusion pores by locking them into the thin catenoidal fusion pore family.

## Results

### Mathematical model of the fusion pore

During exocytosis, content molecules are released through the fusion pore, the initial connection between the fused vesicle and cell membrane. We consider a somewhat simpler geometry, a fusion pore between two planar membranes of diameters 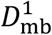 and 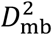, separated by distance *h* maintained by a force *f* (Fig. 1A). The classic Helfrich free energy (39) *F* includes contributions from bending *F*_bend_ and tension *F*_tension_,

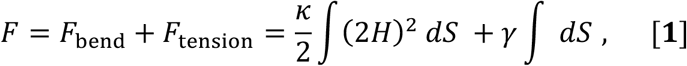

where *κ* is the bending modulus, *H* = (*c*_1_ + *c*_2_)/2 is the mean curvature, the average of the two principle curvatures *c*_1_ and *c*_2_, *γ* is the membrane tension, and the integral is over the membrane surface *S*. We take *κ* = 20 kT (51).

**Figure 1.**
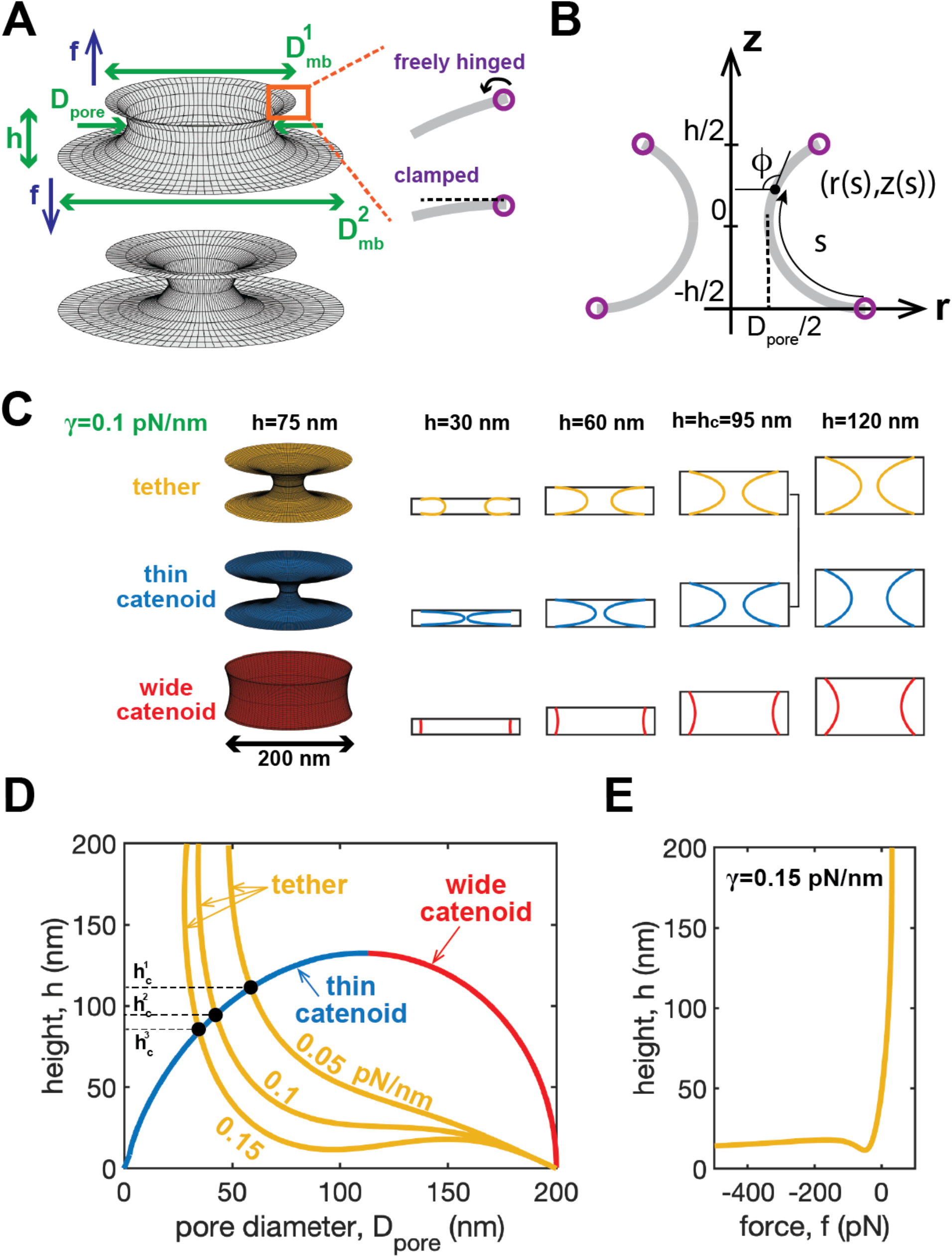
Three families of fusion pores between two planar membranes. **(A)** We study fusion pores between two planar membranes of diameters 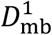 and 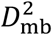 with a separation *h* maintained by a force *f*. Two boundary conditions (BCs) are considered, with and without bending constraints from planar membranes (clamped and freely hinged, respectively). Freely hinged BCs describe soap films, but proper treatment of microscopic membranes requires clamped BCs. The fusion pore diameter *D*_pore_ is the minimal cross-sectional diameter. **(B)** Coordinate system of an axisymmetric fusion pore. **(C)** The three fusion pore families for freely hinged BCs. Examples of the wide catenoid, thin catenoid and tether family are shown for *D*_mb_ = 200 nm and membrane tension *γ* = 0.1 pN/nm. At the critical separation *h_c_*~95 nm, the tether and thin catenoid are identical. **(D)** Pore diameter vs membrane separation for the three families (freely hinged BCs). The two catenoids do not depend on membrane tension, while the tether becomes thinner at higher tension. For high tensions the tether diameter depends non-monotonically on membrane separation. **(E)** For high tensions the tether requires a non-monotonic compressive force to maintain the pore height.

An equilibrium fusion pore shape ***<u>X</u>*** satisfies *δF*(***<u>X</u>***)/*δ**<u>X</u>*** = 0, which leads to the shape equation (see refs. (49, 52)),

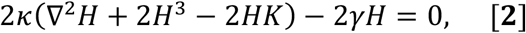

where ∇^2^ is the Laplace–Beltrami operator on the surface and *K* is the Gaussian curvature defined as the product of the two principle curvatures.

Assuming an axisymmetric fusion pore shape for simplicity, with the vertical axis *z* being the symmetric axis (Fig. 1B), the fusion pore is a surface generated by revolution of a curve in the meridian plane. Parameterizing this contour curve by its arclength *s*, the shape equation Eq. **2** is then expressed as an ordinary differential equation involving the vertical coordinate *z*(*s*), the cross-sectional radius *r*(*s*), and the tangent angle *ϕ*(*s*) (48) (Fig. 1B, and see SI Text for a derivation),

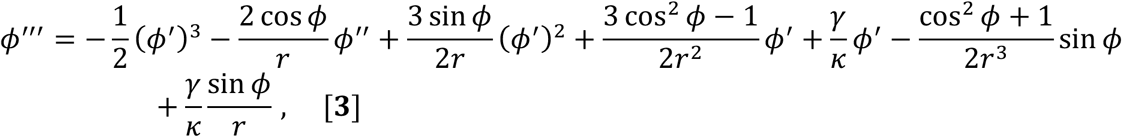

where prime denotes derivative with respect to *s*. This is the same shape equation as for the tether analysis of ref. (48), but our boundary conditions (BCs) are different: the membrane tangent points inwards at the outer edge of both upper and lower membranes, *dr*/*ds* > 0 at *z* = *h*/2 and *dr*/*ds* < 0 at *z* = −*h*/2 (Fig. 1B). In ref. (48) the upper membrane tangent points outwards at the edge.

### Three fusion pore families: wide catenoid, thin catenoid, tether

A proper treatment must account for the bending constraints from the upper and lower planar membranes. However, it is helpful to first study solutions to the shape equation, Eq. **3**, for the simplified case of freely hinged BCs (Fig. 1A, zero torque, or equivalently *H* = 0) at the boundary of the planar membranes. As shown previously (45, 53), there are three fusion pore solutions: two catenoids and one tether-like solution. We consider first equal membranes, 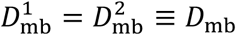. The stability of the three solutions will be discussed in later sections.

#### Wide and thin catenoids

The freely hinged BCs allow for two catenoidal solutions (45, 49). Consider first a fusion pore that is a minimal surface (54), locally minimizing the membrane area *A* and the tension contribution to the free energy of Eq. **1**, *δ*^(1)^*F*_tension_ = *γδ*^(1)^*A* = 0. Parameterizing the surface ***<u>X</u>***(*u*, *v*) with coordinates (*u*, *v*), the surface area element *dS* = *g*^1/2^*dudv*. Thus, a minimal surface satisfies (55) (see SI Text for a derivation)

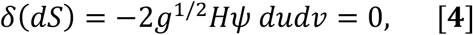

for arbitrary variation *δ**<u>X</u>***. Here *g* is the determinant of the metric tensor and *ψ* is the normal component of *δ**<u>X</u>*** (the tangential component reparametrizes the surface without affecting *A*). Thus, a minimal surface has zero mean curvature, *H* (54). Expressing *H* in terms of our coordinate system *r*(*z*) for an axisymmetric surface and solving *H* = 0 leads to the general catenoidal solution (49),

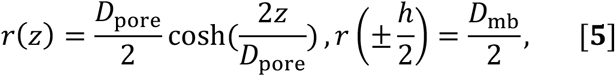

where *D*_poae_ is the pore diameter, the minimal cross-sectional diameter (Fig. 1A). For a given pore height *h*, two values of *D*_poae_ satisfy the BCs of Eq. **5**: 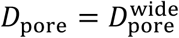 (wide catenoid) and 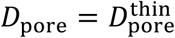 (thin catenoid), see Fig. 1.

Since the mean curvature *H* = 0, the catenoid simultaneously minimizes the positive definite bending free energy, *κ*/2 *f*(2*H*)^2^ *dS*, and satisfies the freely hinged BCs. Thus, both catenoids satisfy *δ*^(1)^*F* = 0 (Eq. **1**). Since the catenoids minimize the area regardless of membrane tension *γ*, the catenoidal solutions are unaffected by the value of *γ*. Note the freely hinged BC (*H* = 0) guarantees an exact catenoid solution as the BC cost no additional bending energy.

#### Tether

Numerical solution of Eq. **3** yields a third “tether” solution family (45) in addition to the two catenoids. For illustration Fig. 1C shows tethers for *D*_mb_ = 200 nm (similar to the size of a dense-core vesicle (56, 57)) and membrane tension *γ* = 0.1 pN/nm (58). For large separations, the third tether solution adopts almost cylindrical shapes whose radius equals the capillary length *λ* ≡ (*κ*/2*γ*)^1/2^, similar to a tether pulled from a membrane (48, 49). For small separations, tethers require a compressing force (*f* < 0) between the two planar membranes, unlike catenoids (Fig. S1). In contrast to the two catenoids, the shape of the tether solution is tension-dependent (Fig. 1D). For small tensions, the tether has monotonically decreasing pore diameter with increasing membrane separation. However, above a certain tension, neither the pore diameter nor the compressing force *f* are monotonic (Figs. 1D, 1E, S2). Notably in studies of tether pulling (48, 49), an oscillation in the force-height plane was also found for larger *h* values where a cylindrical tether starts to form, but unlike our study those tethers are under a positive pulling force. Regardless of membrane tension, the tether branch always intersects the thin catenoid branch at a critical separation *h_c_* where the two branches share the same shape (Fig. 1C, D). The critical separation decreases with membrane tension.

### The thin catenoid is stable on nanoscales

Thus far we have used the shape equation, Eq. **3**, to determine equilibrium fusion pore shapes. However, whether an equilibrium shape can be observed depends on its stability. Only the fusion pore shapes that locally minimize the free energy *F* can be realized. Thus, we next considered the second variation of the Helfrich energy *δ*^(2)^*F* of each catenoidal surface to determine whether it is positive definite.

#### Soap films: wide catenoids

For macroscopic soap films, the bending energy is a negligible contribution, so *F* ≈ *γ ∫ dS* in Eq. **1** (59). Thus, stability of a soap film depends only on the second variation of the film area *A*, *δ*^(2)^*A*. Under an arbitrary deformation *δ**<u>X</u>*** whose normal component is *ψ*, the second variation can be expressed as (60)

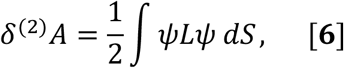

where the operator *L* ≡ −∇^2^ + 2*K* and the integral is over the reference surface (in our case the exact catenoidal surface, Eq. **5**). We consider the eigenvalue problem 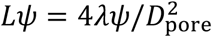, which becomes a Sturm-Liouville problem when we parameterize the axisymmetric deformation *ψ*(*v*) by the normalized vertical coordinate *v* ≡ 2*z*/*D*_poae_ (Fig. 1B, see SI Text for a derivation) (50),

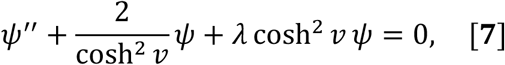

where prime denotes derivative with respect to *v* and the boundary conditions are *ψ*(±*h*/*D*_pore_) = 0. Thus, any arbitrary deformation *ψ* can be expressed in terms of the eigenfunctions *ψ*_i_ (*i* = 1,2,…) of Eq. **7**, *ψ* = ∑_*i*_ *c*_i_*ψ*_i_. (*c*_i_ are constants), and the second variation in the area is 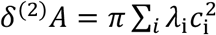 where *λ*_1_ < *λ*_2_ <… are the eigenvalues of Eq. **7**. Thus, whether a deformation mode *ψ*_i_ contributes negatively or positively to *δ*^(2)^*A* depends on the sign of the corresponding eigenvalue *λ_i_*. Note that the thin and wide catenoids have distinct sets of eigenvalues and eigenfunctions determined by the ratio *D*_pore_/*h*.

Numerically solving the eigenvalues and the eigenfunctions of Eq. **7**, it can be shown that for wide catenoids all the eigenvalues are positive, while for thin catenoids one of the eigenvalues is negative, *λ*_1_ < 0 (50) (Fig. S3). Thus *δ*^(2)^*A* is always positive for wide catenoids and they can stably exist, as observed in soap film experiments (61). In contrast, thin catenoids are unstable as the first deformation mode *ψ*_1_ makes *δ*^(2)^*A* negative.

#### Cellular membranes: wide and thin catenoids

On cellular scales (nm to μm) bending energy is important, so the second variation *δ*^(2)^*F* of the full Helfrich energy Eq. **1** must be considered. For a catenoid whose mean curvature *H* is zero, *δ*^(2)^*F* given a normal deformation *ψ* can be greatly simplified to (60) (see SI text)

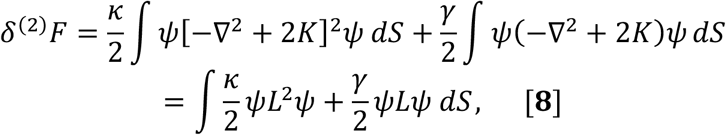

where the integral is over the catenoidal surface. Thus, using the same eigenvalues *λ_i_* and eigenfunctions *ψ_i_* of Eq. **7**, we obtain (see SI Text for derivation)

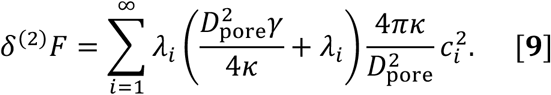

Any deformation mode *ψ*_i_. will destabilize the catenoid if 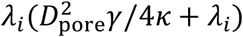 is negative. Consider positive membrane tension *γ*. Then wide catenoids are always stable, as all eigenvalues are positive, but since a thin catenoid has one negative eigenvalue, *λ*_1_ < 0, it is stable only if its spatial dimensions are sufficiently small, 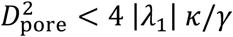, or equivalently *h* < *h_c_* where *h_c_* is the critical membrane separation corresponding to this upper limit for pore size. These stable and unstable regions are shown in Fig. 2D. Note that the limit of stability *h* = *h_c_* on the small catenoid solution is precisely the point where the curve intersects the tether solution, Figs. 1D, 2A. Thus, increasing membrane separation beyond the critical point, *h* > *h_c_* (at fixed tension), the small catenoid becomes unstable and is replaced by the tether. This is an example of a bifurcation occurring when a solution becomes unstable, qualitatively similar to the behavior of a spherical vesicle under pressure (62): above a critical pressure, the spherical solution becomes unstable and two new solution branches appear, prolate and oblate. The stable prolate shape replaces the sphere.

**Figure 2.**
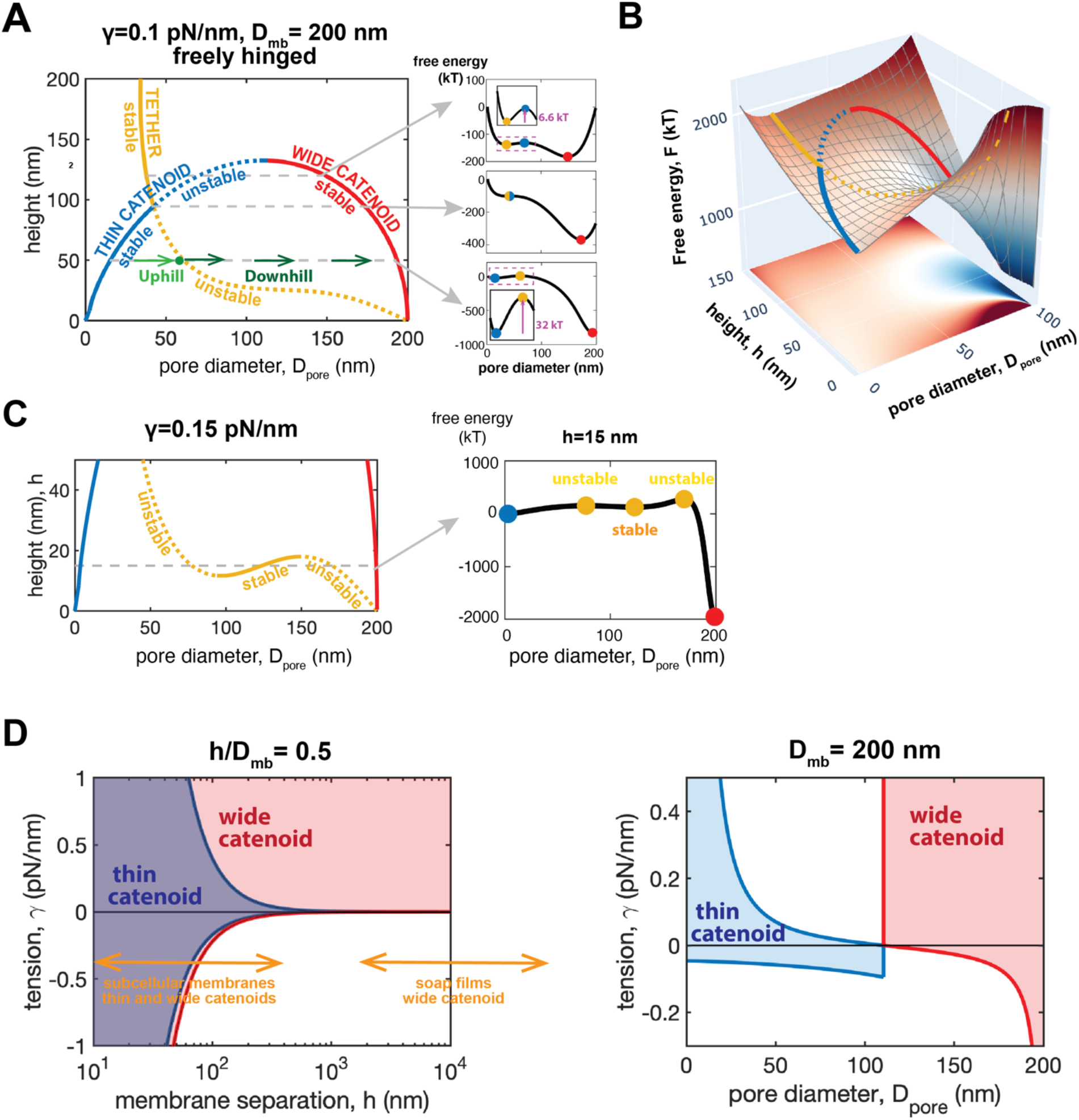
Two microscopic catenoidal fusion pores are observable between freely hinged membranes. **(A)** Stability of the three fusion pore families. At sufficiently small membrane separation, thin and wide catenoids are both stable. Free energy landscape is shown for three h values (right). The stable (unstable) fusion pores localize in valleys (ridges). Pore dilation from a thin to wide catenoid requires surmounting an energy barrier set by the tether. **(B)** Free energy landscape. **(C)** For higher membrane tensions and sufficiently close membrane there are five pore families, three of which are stable. **(D)** Thin catenoidal fusion pores are only observable on microscopic scales within band of tensions which shrinks to zero on macroscopic scales.

Stability can also be phrased in terms of tension, *γ*. For wide catenoids, the lowest eigenvalue *γ*_1_ sets a negative lower bound on the tensions for which stability holds. For thin catenoids the negative lowest eigenvalue and lowest positive eigenvalue, *λ*_1_ and *λ*_2_, set an upper and lower bound on tension, respectively. Thus

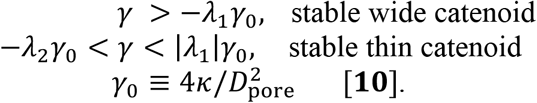

Since the characteristic tension *γ*_0_ decreases with increasing spatial dimension (measured by *D*_pore_), it follows that thin catenoids are observed only at small scales: as the scale increases the band of tensions where thin catenoids are stable shrinks and tends to zero, Fig. 2D. Thus, at macroscopic scales (soap films) only wide catenoids are seen. Thin catenoids are specific to scales found in cells. For example *γ*_0_~10^-1^ pN/nm for *D*_pore_~50 nm and *κ* =20 kT (51), comparable to cellular membrane tensions (63). These results are consistent with ref. (45) where the buckling transitions of fixed-area catenoidal membranes were studied.

### Free energy landscape for fusion pore expansion

The three families (thin catenoid, wide catenoid and tether) are different minimum energy fusion pore shapes with different fusion pore diameters, *D*_pore_. Each family defines a valley (or ridge, if unstable) in (say) the *h* – *D*_pore_ plane (Fig. 1D). What free energy landscape do these valleys belong to? What barriers separate the valleys?

To address this we solved the shape equation, Eq. **3**, constraining the pore diameter to have an arbitrary value *D*_pore_ (see SI Text). The radial force to maintain a pore with this diameter is 2*κ*(*dH*/*ds*) evaluated at the pore (64). Integrating this force over a range of *D*_pore_ values and using the free energies of thin catenoids calculated from Eq. **1** as the reference levels, we obtained the free energy landscape of Figs. 2A-C.

For a given membrane separation *h*, the free energy reaches maxima or minima at the pore diameters corresponding to the three solution families (Fig. 2A). Consistent with the stability analysis, the thin catenoid where predicted to be stable (unstable) lies in a valley (on a ridge). The tether solution has the opposite stability to the thin catenoid. For any separation *h*, the wide catenoid has the lowest energy, but there is an energy barrier Δ*F* opposing the fusion pore from expanding from a thin to wide catenoid (Fig. 2A), whose height equals the free energy difference between the thin catenoid and the tether solution. The exception is the critical separation *h_c_* where the thin catenoid merges with the tether and the energy barrier vanishes (Fig. 2A).

For high tensions, the tether solution oscillates in the *h* – *D*_poae_ plane at small separations, with five solutions (two catenoids and three tethers) for a given separation (Fig. 2C). Calculating the free energy landscape as a function of *D*_poae_ at such a fixed *h* we found three valleys and two ridges as the fusion pore expands. Thus, in addition to the thin catenoid, a second metastable fusion pore is the tether with intermediate pore diameter in the middle valley.

### With realistic membrane bending constraints there are three fusion pore families

Next we solved the shape equation, Eq. **3**, accounting properly for the bending constraints supplied by the planar membranes connected by the fusion pore. We used the appropriate zero slope BC at the boundaries, not freely hinged BCs. The exact catenoids are no longer solutions as they violate the boundary conditions, but we find there are still three fusion pore families (Figs. 3A, S4).

**Figure 3.**
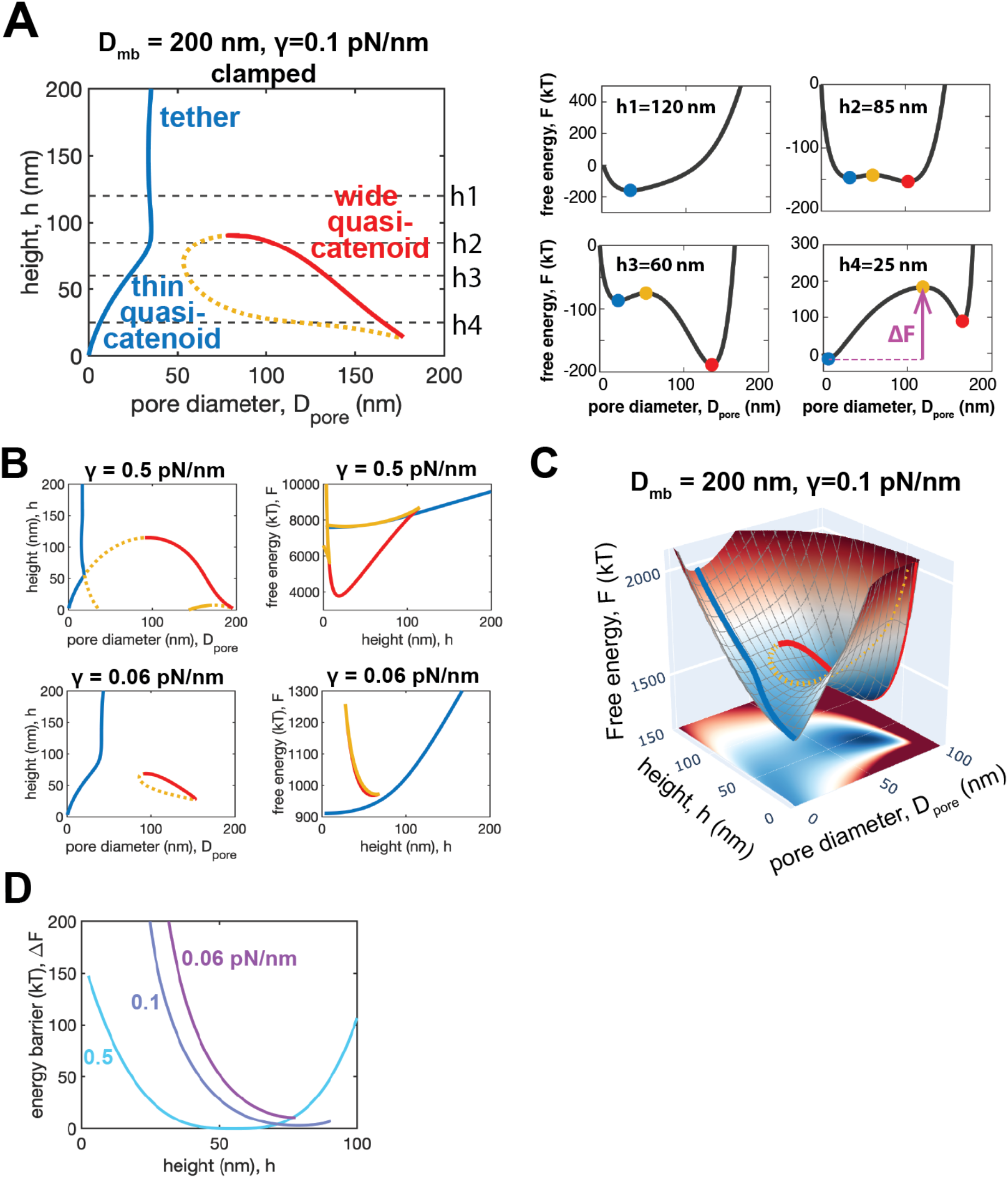
A narrow and a wide fusion pore family are stable for realistic membrane bending constraints. Solid lines: stable. Dashed lines: unstable. **(A)** Stability of the three fusion pore families with realistic clamped boundary conditions. Free energy landscape for four values of *h* (right). Dilation from the thin to the wide catenoidal pores requires crossing an energy barrier Δ*F* set by the third unstable solution. **(B)** Three fusion pore families and their free energies for two membrane tensions. **(C)** Free energy landscape. **(D)** Energy barrier for pore dilation vs height for three membrane tensions. The barrier is lower at higher tensions.

Again, we find there are three pore families whose properties depend on the capillary length *λ* ≡ (*κ*/2*γ*)^1/2^. Scales smaller (larger) than *λ* are dominated by bending (membrane tension). For high membrane tensions such that the capillary length is smaller than the membrane diameter *D*_mb_, the three families are close to those freely hinged BCs (Figs. 3A, 3B, S4). For physiologically relevant small separations, the fusion pore with the smallest width resembles the thin catenoid (we call it the thin quasi-catenoid) while the pore with the largest width resembles the wide catenoid (we call it the wide quasi-catenoid). A third unstable family with intermediate width appears. Note that switching on the bending constraint disconnects the quasi-thin catenoid and the unstable pore families.

### Realistic membrane bending constraints: free energy landscape

Accounting for the bending constraints provided by the planar membranes (clamped BCs), we repeated the free energy calculations previously performed for the simplifying case of freely hinged BCs (Figs. 2A-C). We find a qualitatively similar landscape (Figs. 3A, 3C). Again, an energy barrier Δ*F* separates the locally stable thin and wide quasi-catenoids whose height is defined by the unstable intermediate branch (Fig. 3A). For example, for membrane separation 60 nm and membrane tension 0.1 pN/nm, the barrier is ~10 kT. To dilate the pore this barrier must be crossed.

The free energy landscape is highly sensitive to membrane tension (Fig. 3B). Increased tension lowers the free energy barrier Δ*F* for physiologically relevant small heights (Fig. 3D). When tension is below a certain threshold, the wide quasi-catenoid family disappears (Fig. S5), so that dilation of the narrow thin catenoid is no longer possible. These findings are consistent with experiments showing high membrane tension favors full-fusion exocytosis (24) and correlates with larger fusion pores (23).

### Fusion pore between a nanodisc and a planar membrane

Membrane nanodiscs (NDs) are powerful systems for the study of individual pores (29–31, 65, 66). NDs are nanosized phospholipid membrane patches bounded by a lipoprotein scaffold forming a wall of two parallel alpha helices (67). By stabilizing fusion pores, NDs allow pore sizes and fluctuations to be inferred from conductance measurements. However, the mechanism of pore stabilization and the relation to pores in larger membranes of phycological relevance is not understood.

Here we used a simple elastic model (31) to determine the torque required to rotate the scaffold a certain angle and we repeated pore shape calculations with this torque-balance BC at the ND edge (Fig. 4A, see SI text). This BC is intermediate between the clamped and freely hinged limits. With a typical ND diameter *D*_ND_ = 24 nm and membrane tension 0.1 pN/nm, we find the ND fusion pores can only adopt thin quasi-catenoidal shapes (Figs. 4B, C). The wide quasi-catenoid is theoretically also accessible (Fig. 4D), but the membrane is forced to bend outward at the ND edge, rotating the scaffold > 90°. We suggest this would disassemble the scaffold and the ND. Furthermore, an energy barrier > 150 kT(set by the third unstable pore solution) impedes transition to this wide pore (Fig. S7). Our results also suggest the ND studies are not able to probe fusion pores under high membrane tension, as tensions as high as 2 pN/nm may activate the wide catenoidal pore since the barrier is lowered to ~10 kT (Fig. S7). In summary, we conclude that NDs stabilize fusion pores by banishing access to the dilated wide catenoidal pore.

**Figure 4.**
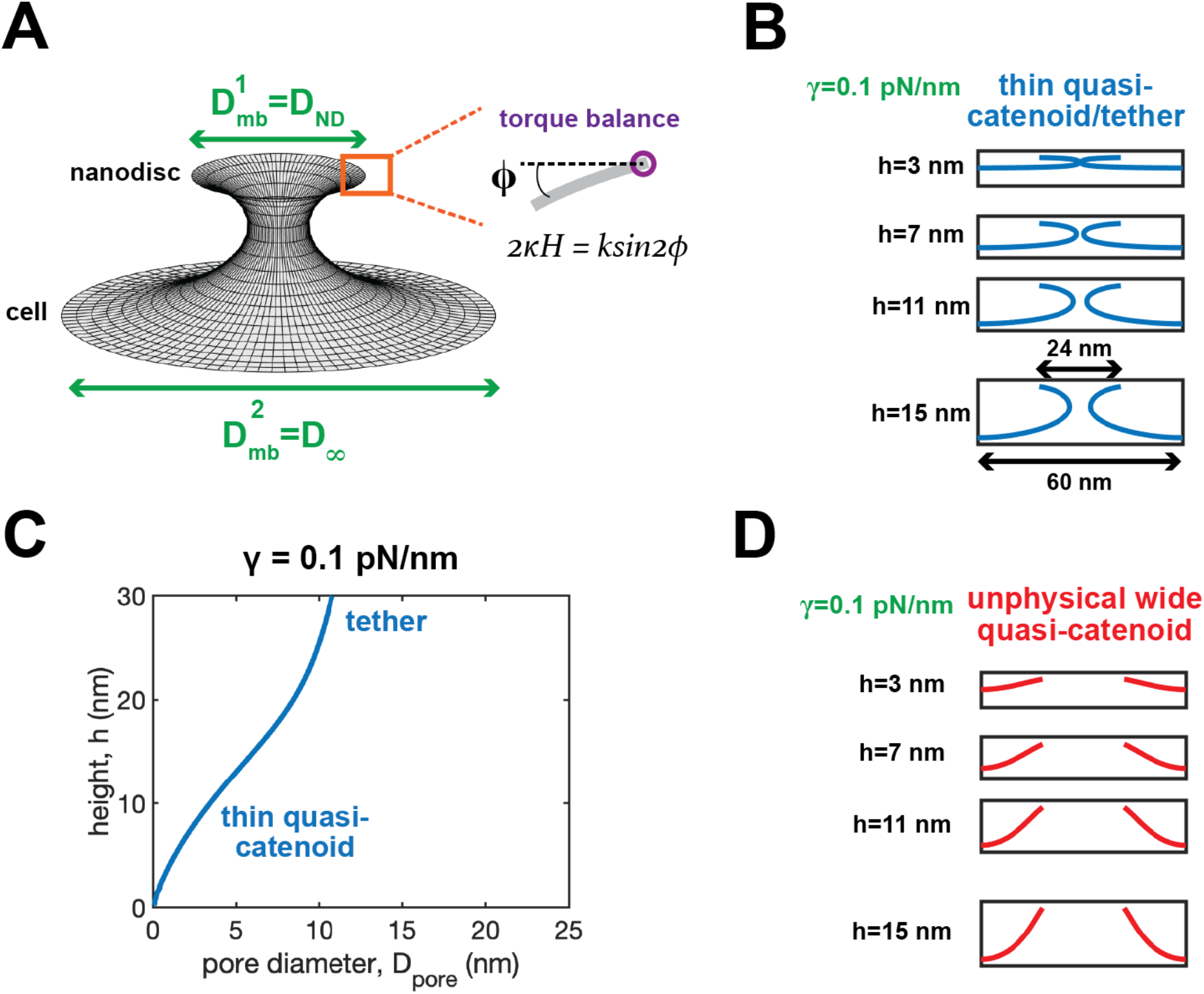
Nanodiscs lock fusion pores into thin quasi-catenoids. **(A)** Fusion pore between a nanodisc (ND) and a planar membrane. A torque balance boundary condition is imposed at the upper membrane edge, representing resistance to twisting from the ND scaffold. **(B)** Thin quasi-catenoid fusion pores. **(C)** Pore diameter vs height for the thin quasi-catenoid pore family. The thin quasi-catenoid develops into tether-like shapes for large *h*. **(D)** Unphysical wide quasi-catenoid pores. The ND scaffold is twisted by a large angle *ϕ* > 90° which may disassemble the scaffold.

## Discussion

Membrane fusion pores are critical to many fundamental processes such as neurotransmission, hormone release, trafficking and fertilization (10). Here we identified three fusion pore families: two quasi-catenoids and a third unstable family that sets the scale of the energies (Fig. 3A). The thin catenoid family has the narrowest waist, and is unique to the microscopic scales of cellular membranes, being stabilized by membrane bending forces, while the wide catenoid is a fully dilated pore. These conclusions are based on properly accounting for the bending constraints provided by the membranes away from the fusion pore. For sufficiently low membrane tensions or large membrane separations, the wide catenoid fusion pore no longer exists, leaving only thin catenoids (Figs. 3A, B). We call the pore of intermediate size the unstable intermediate; however, for certain membrane separations this family in fact has regions of stability, corresponding to oscillations in the membrane separation-pore size plane (*h* – *D*_pore_ plane of Fig. 3B). A related oscillation was found for pulled membrane tethers (48).

In contrast to microscopic membrane surfaces, macroscopic soap films adopt only the wide catenoid, as bending forces contribute negligibly so that thin catenoids are always unstable. Indeed, using the freely hinged BCs appropriate to soap films, we showed the thin catenoid family is then stable for a band of tensions (with positive and negative upper and lower bounds) which shrinks to almost zero for macroscopic scales, while the wide catenoid is stable for any tension above a minimum value which tends to zero on macroscopic scales (Fig. 2D).

During exocytosis vesicle-plasma membrane fusion generates a small nascent ~1 nm fusion pore (19, 27), which must dilate if full release of the vesicle contents is to occur. Pore dilation has been characterized by amperometry recordings during dense core vesicle release, showing an initial small flickering pore (prespike foot) that suddenly dilates to a large pore (spike) (18–20, 68). The energy barrier for membrane fusion was estimated (69), but the energy landscape for pore dilation has not been clear. Here, the three families of fusion pores from our analysis suggest the initial flickering pore belongs to the stable narrow pore family, and the abrupt dilation is a transition to the stable wide pore family over a barrier whose height is set by the unstable intermediate pore family (Fig. 5). For example, for ~ 400 nm dense core vesicles in chromaffin cells, we estimate the barrier is ~50 kT, depending on membrane tension and separation (Fig. 3D). This energy landscape that the three pore families belong to is the landscape that pore size-regulating factors such as dynamin, amisyn, Syt and SNARE family proteins (21, 29–31, 35) must navigate (Fig. 3C). Its knowledge will help to establish the molecular mechanisms controlling pore dilation.

**Figure 5.**
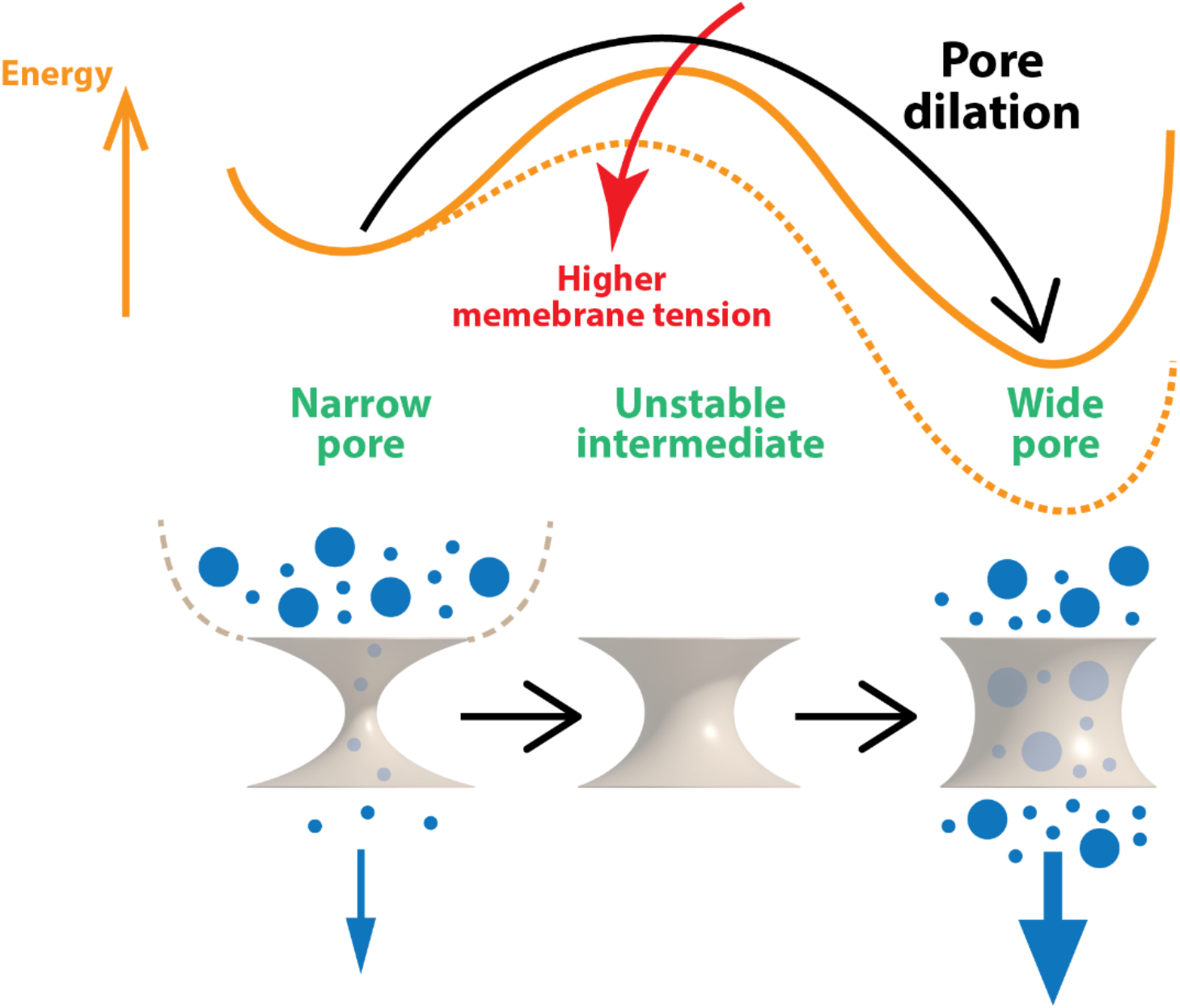
Model of the pore dilation transition during exocytosis. Three fusion pore families define the dilation pathway. Membrane fusion generates a narrow pore permitting release of smaller molecules only. Dilation to the wide pore family requires a barrier to be surmounted, whose height is set by the unstable intermediate pore and is lowered by increased membrane tension. The wide pore permits full contents release including larger molecules.

An important study showed that the initial small fusion pore is longer lived for larger vesicles, reflected by amperometric feet of greater duration (70). It was argued that larger vesicles have dilated pores of higher energy, explaining the increased dilation delay time. Correspondingly, we also found that the dilated (wide quasi-catenoidal) pore has higher energy for larger fusing membranes (larger *D*_mb_, Fig. S6). However, more importantly the free energy of the initial (small quasi-catenoidal) pore increased by a greater amount, primarily because its area increased by a greater amount. The unstable intermediate pore energy increased only moderately. Overall, the barrier to dilation decreased for larger vesicles. We speculate that other factors such as the mechanisms of SNARE- and synaptotagmin-mediated pore expansion (29–31) may depend on vesicle size.

Membrane tension has been shown to correlate with larger fusion pores and favor full-fusion over kiss-and-run (23, 24). Our analysis rationalizes these observations, as we found the barrier to pore dilation decreases with increasing membrane tension (Figs. 5, 3D). As membrane tension decreases, dilation becomes increasingly energetically demanding, until below a certain tension dilation cannot occur as the stable wide pore family ceases to exist (Fig. S5).

Finally, we studied fusion pores between a nanodisc and a large planar membrane (such as an extended cell plasma membrane). Nanodiscs are nanoscale membranes, powerful systems to study single fusion pores by conductance measurements (29–31, 65, 71), but it has been unclear how representative these fusion pores are of physiological pores. For typical 10-30 nm nanodiscs, we find only the narrow pore family is accessible. Theoretically, if a large energy barrier is surmounted wide pores are also accessible. However, the wide catenoid requires outward-oriented membrane at the nanodisc edge (Fig. 4D), a disruption imposing large twisting forces that we suggest would destabilize the nanodisc. In effect, only the small nascent fusion pore is realizable. This quantifies how nanodiscs stabilize fusion pores, by locking them into the narrow pore family and blocking dilation.

## Acknowledgements

This work was supported by National Institute of General Medical Sciences of the National Institutes of Health under award number R01GM117046. The content is solely the responsibility of the authors and does not necessarily represent the official views of the National Institutes of Health. We acknowledge computing resources from Columbia University’s Shared Research Computing Facility project.

## Author contributions

B.O’S designed the research. R.S and B.O’S performed mathematical analysis. R.S, S.W and Z.M performed numerical calculations. B.O’S, R.S wrote the paper with contributions from Z.M.

## Supporting Information

### Model description

Here we present a concise derivation of the shape of catenoid, a minimal surface, and the shape equation for axisymmetric membranes, the stability analysis of catenoidal membranes with subject to classic Helfrich free energy (Eq. 1, (1)) using variational principles.

#### Basic differential geometry

We parameterize an arbitrary surface ***<u>X</u>***(*u*^1^, *u*^2^) with coordinate (*u*^1^, *u*^2^). The two tangent vectors *e*_1_, *e*_2_ are

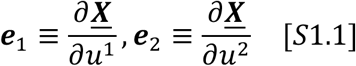

The metric tensor, or the first fundamental form, is defined as

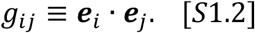

The inverse metric *g^ij^* is the inverse of the metric tensor *g_ij_* such that

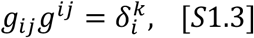

where 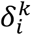 is the Kronecker symbol. The second fundamental form *b_ij_* is given by

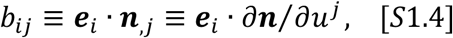

where ***n*** is the normal vector given by

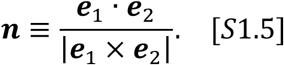

The mixed second fundamental form 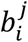 is defined as

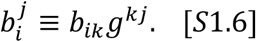

The mean curvature *H* is defined as

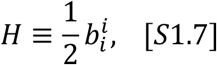

while the Gaussian curvature G is defined as

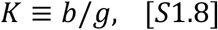

where *g* and *b* are the determinant of the first fundamental *g_ij_* and the second fundamental *b_ij_*.

#### Minimal surface

A minimal surface is a surface whose area is locally minimized. To derive a minimal surface we consider the area variation with subject to an infinitesimal perturbation ***<u>X</u>*** → ***<u>X</u>*** + *δ**<u>X</u>***. The perturbation *δ**<u>X</u>*** has a normal component (in the direction of ***<u>n</u>***) and two tangent components (in the directions of ***<u>e</u>***_1_ and ***<u>e</u>***_2_),

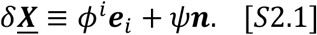

The variation in the tangent vector is

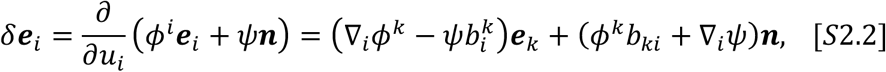

where ∇_*i*_ is the covariant derivative with respect to *u_i_*. Note that *δ**e**_i_* only has the first order term. The variation of the first fundamental form *g_ij_* has a first order term *δ*^(1)^*g_ij_* and a second order term *δ*^(2)^*g_ij_*,

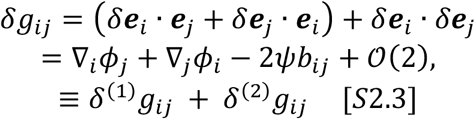

where *ϕ_i_* ≡ *ϕ^j^g_ji_*. The variation of the determinant *g* is then

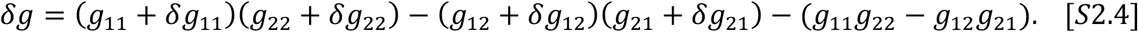

Extracting the first order term *δ*^(1)^*g* in *δg*, we have

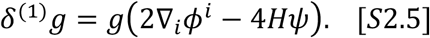

The infinitesimal area 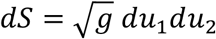. The first variation in area *δ*^(1)^*A* is

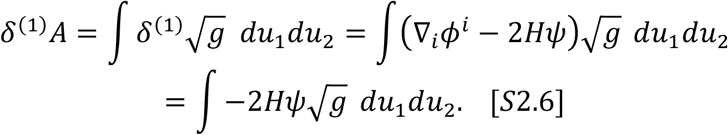

Note since the tangent terms only reparametrize the surface and do not change the area, we only keep the normal term.

A minimal surface requires *δ*^(1)^*A* = 0. Thus a minimal surface has zero mean curvature *H* = 0.

#### Catenoid

To derive the shape of catenoid, the axisymmetric minimal surface, we parametrize the surface by the arclength of the contour in a meridian plane *s* and the azimuthal angle *θ*, such that

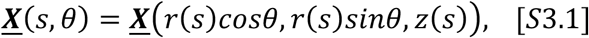

where *r* and *z* is the correctional and vertical coordinate, respectively (Fig.1B). The two tangent vectors are

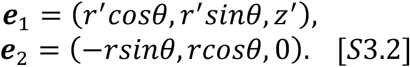

The first fundamental *g_ij_* is

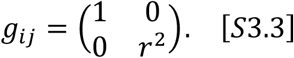

The second fundamental *b_ij_* is

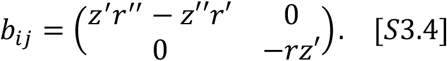

The mean curvature *H* is

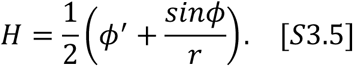

where *ϕ* is the tangent angle (Fig. 1B, different from the tangent variation *ϕ^i^*) such that *tanϕ* = –*dz*/*dr*. To solve the shape of catenoids *r*_cat_(*z*), we set *H* = 0 which gives

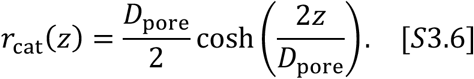

#### Shape equation

The shape of the membrane that minimizes the classic Helfrich free energy (Eq. 1) has to satisfy the shape equation

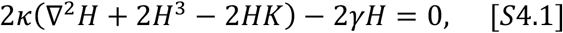

For an axisymmetric membrane the Laplace–Beltrami operator ∇^2^ is

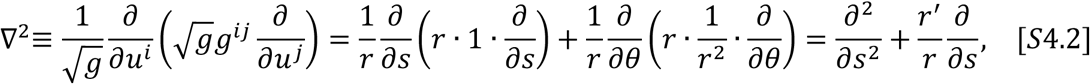

Note that *∂*/*∂∂* = 0 for axisymmetric membranes. Also the Gaussian curvature *K* is

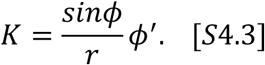

By plugging Eq. S3.4, Eq. S4.3 into Eq. S4.2 we derive the shape equation for axisymmetric membranes (Eq. 3). Note that *r*′ = *cosϕ*.

#### Catenoid stability

To analytically derive the stability of the catenoidal membranes, we consider the second variation. For soap films with no bending energy, we consider the second variation in the area, *δ*^(2)^*A*. For a membrane of an arbitrary shape,

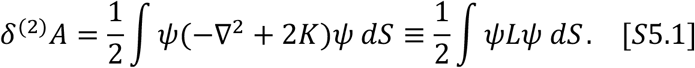

The derivation is similar to that of the first variation *δ*^(2)^*A*, but the variations in the metric tensor elements *g_ij_* have to be expanded to second order. A detailed derivation can be found in ref. (2).

We parameterize the catenoidal surface *S_cat_* with two parameters, azimuthal angle *u* and normalized height *v*, such that

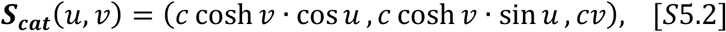

where *c* is half of the pore diameter *D*_pore_ (Fig. 1A).

The two tangent vectors are

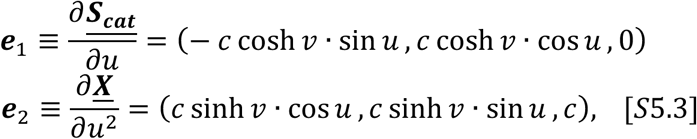

The metric tensor *g_ij_* is

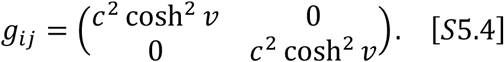

The determinant *g* is

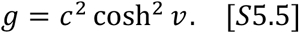

The Laplace–Beltrami operator ∂^2^ is

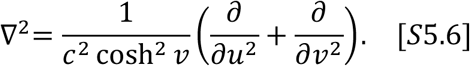

The Gaussian curvature *K* is

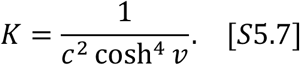

Thus keeping only the axisymmetric term *∂/∂v*^2^ the operator *L* = −∇^2^ + 2*K* is

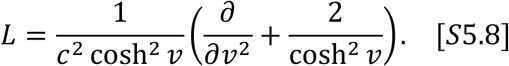

The eigenvalue problem *Lψ* = *λψ*/*c*^2^ leads to a Sturm-Liouville problem

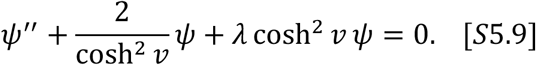

whose eigenfunctions and eigenvalues are *ψ_i_* and *λ_i_*. For a perturbation 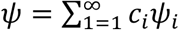 Thus, in terms of *ψ_i_* and *λ_i_* the second variation in area is

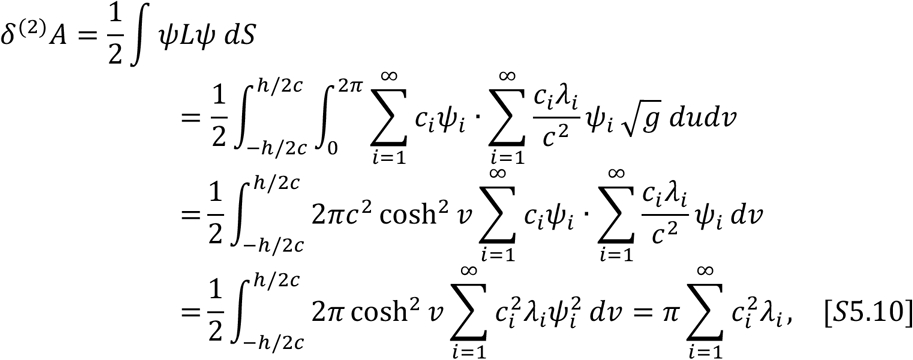

where *h* is the height of the catenoid. The properties of Sturm-Liouville eigenfunctions are used

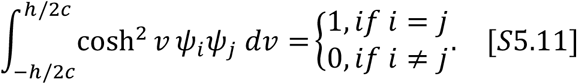

With the inclusion of bending energy, the second variation in Helfrich free energy (Eq. 1) is

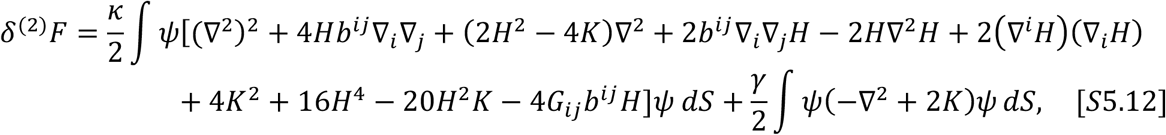

where *b^ij^* is the inverse of the second fundamental form *b_ij_*, ∇^*i*^ ≡ *g^ij^*∇_*j*_, *G_ij_* is the Einstein tensor and the integral is over the reference surface. A detailed derivation can be found in ref. (2)

For a catenoid whose mean curvature is zero, Eq. S5.9 can be greatly simplified to

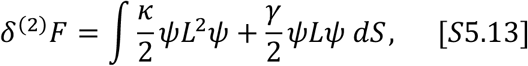

The variation in the bending term *δ*^(2)^*F*_bend_ is

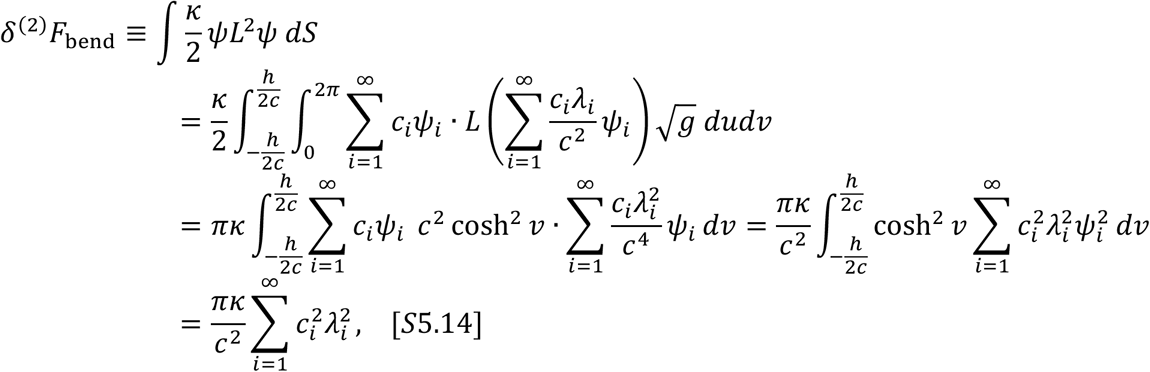

Thus the second variation of the free energy is

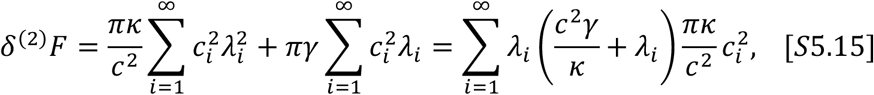

#### Radial force on the pore

To calculate the free energy landscape of fusion pores with an arbitrary diameter *D*_pore_, we need to calculate the radial force acting on the pore to maintain *D*_pore_. Because the two membranes have the same size, 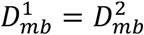 (Fig. 1A), the minimal cross-sectional diameter will always appear in the middle, *z* = 0. Thus, we solve the same shape equation Eq. 3 but only solve the shape of half of the pore (−*h*/2 < *z* < 0) where the imposed pore diameter comes into the boundary condition at *z* = 0. To calculate the radial force acting on *z* = 0 we use the stress tensor (a derivation see ref. (3))

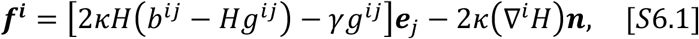

where ***f^i^*** is a 3 × 2 tensor.

The force per unit length exerted on the boundary *z* = 0 is

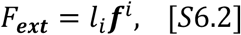

where *l_i_*(*i* = 1,2) are constants such that *l_i_**e_i_*** is the normal vector that lies in the tangent plane and is perpendicular to the boundary. Here *l_i_**e_i_*** is the unit vector pointing in the *z* direction (Fig. 1B). Using the axisymmetric parameterization ***<u>X</u>***(*s*, *ψ*) (Eq. S3.1), the two normal vectors at the boundary *z* = 0 where *ϕ* = *π*/2 are

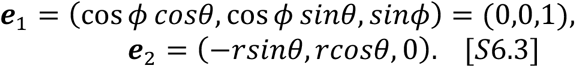

Thus *l*_1_ = 1 and *l*_1_ = 0. *F_**ext**_* is then

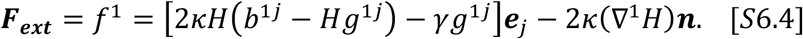

Since we are interested in the radial force we only extract the normal component,

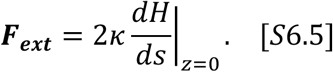

The free energy change Δ*F* from a pore diameter 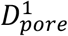 to anther diameter 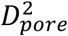 is

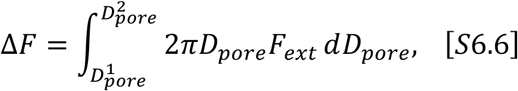

where the factor of 2 comes from the fact that both the upper (0 < *z* < *h*/2) and the lower membrane (−*h*/2 < *z* < 0) are exerting forces on the boundary.

#### Torque balance boundary condition

To more realistically model the nanodisc (ND) fusion pores, we further used a torque-balance boundary condition where the fusion pore meets the ND scaffold (Fig. 4A). Our torque-balance condition relies on a simple elastic model of the ND scaffold originally developed in ref. (4). Because of the two-alpha helix structure of the ND scaffold, it is more difficult to bend the scaffold in one material direction than another. Approximating the cross section of the scaffold as a rectangle with dimensions *w* × 2*w*, where *w* = 1.2 nm is the diameter of a typical alpha helix, it is much harder to bend the scaffold across its wider face (in the transverse direction) than across the thinner face (in the lateral direction).

The elastic energy of the scaffold is given by

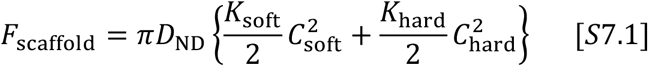

where *D*_ND_ is the ND diameter, *K*_soft_ and *K_hard_* are the respective moduli for lateral and transverse bending modes of the scaffold, and *C*_soft_ and *C*_hard_ are the respective material curvatures of the scaffold in the lateral and transverse directions. Classical elasticity theory shows that, for a rod with a *w* × 2*w* rectangular cross section, the bending moduli are given by

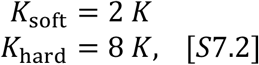

where *K* is the bending modulus of a *w* × *w* rod. The bending modulus of a typical single alpha helix is *K* ≈ 100 *kT* · nm; thus the ND scaffold has bending moduli *K*_soft_ = 200 *kT* · nm, *K*_hard_ = 800 *kT* · nm. We find that torque balance between the membrane and the ND scaffold is achieved when

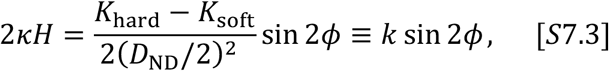

where *ϕ* is the angle by which the cross section of the ND scaffold has been rotated, and *k* is the twisting stiffness of the ND (see SI). A non-elastic soft scaffold (*K*_hard_ = *K*_soft_) will lead to freely hinged boundary condition (*H* = 0) while a stiff scaffold (*K*_hard_ » *K*_soft_) will give a clamped zero-slope boundary condition.

To model the fusion pore of a membrane ND fusing with a cell membrane, we solved the shape equation Eq. **2**, applying the torque balance condition, Eq. **15**, at the ND edge, *z* = *h*/2, and clamped boundary conditions where the fusion pore meets the cell membrane, *z* = –*h*/2. The ND scaffold is assumed to have a diameter of 24 nm, while the diameter of the boundary meeting the cell, *D*_∞_, is chosen to be a much larger value, 60 nm (4) (Fig. 4A).

### Numerical methods

We used MATLAB boundary value problem solver ‘bvp4c’ to solve the shape equation Eq. S8.1 (identical to Eq. 3) to obtain exact shapes of fusion pores.

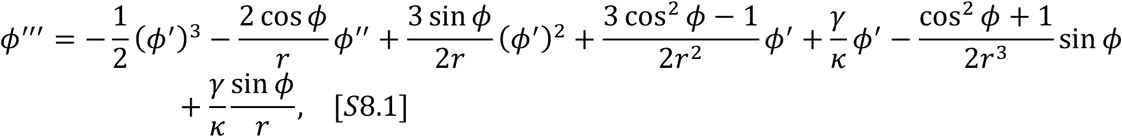

The shape equation is parameter by the arclength *s* and is a third order ordinary differential equation. The radial and vertical coordinates *r*(*s*) and *z*(*s*) is correlated to the tangent angle *ϕ* by

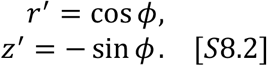

Note that the total arclength *s_max_* is also an unknown parameter. Since bvp4c requires a definite range for the independent variable, here we used the normalized arclength 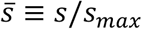 as the independent variable and we solve the shape equation on *s* ∈ [0,1].

Thus we need a total of six boundary conditions (BCs). Each of the boundary, either at 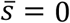 or 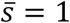, contributes three of them. For a fusion pore with freely hinged BCs, the six BCs are

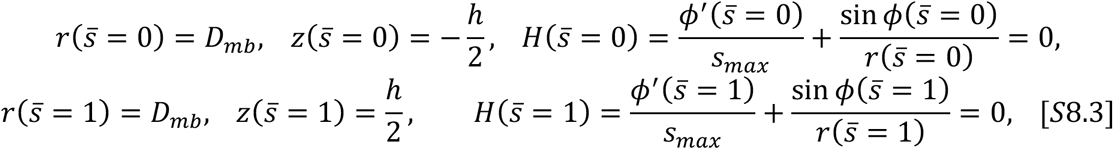

For a fusion pore with clamped BCs, the six BCs are

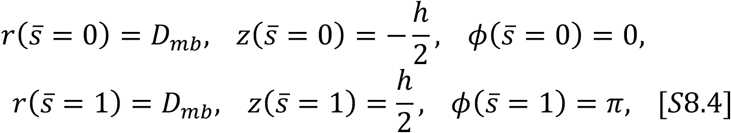

For a ND fusion pore with a torque balance BC, the six BCs are

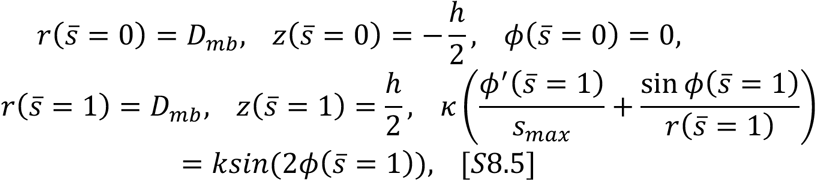

For a pore with an imposed *D*_pore_, the six BCs (for the clamped case) are

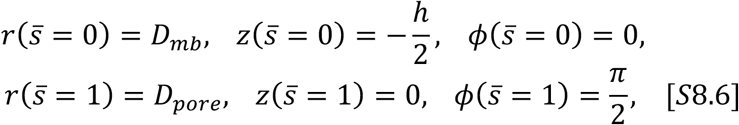

For a pore with an imposed *D*_pore_, the six BCs (for the freely hinged case) are

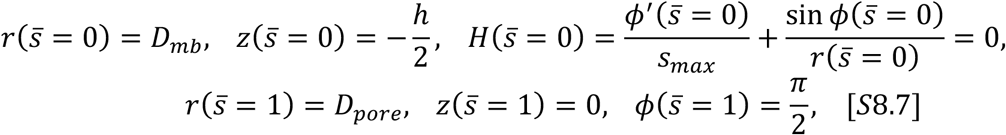

The eigenvalue problem *Lψ* = *λψ*/*c*^2^ is a Sturm-Liouville problem (Eq. S5.9), and we numerically solved it using the MATLAB package MATSLISE (5). The boundary conditions are

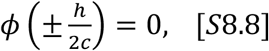

## Supplementary Figures

**Figure S1.**
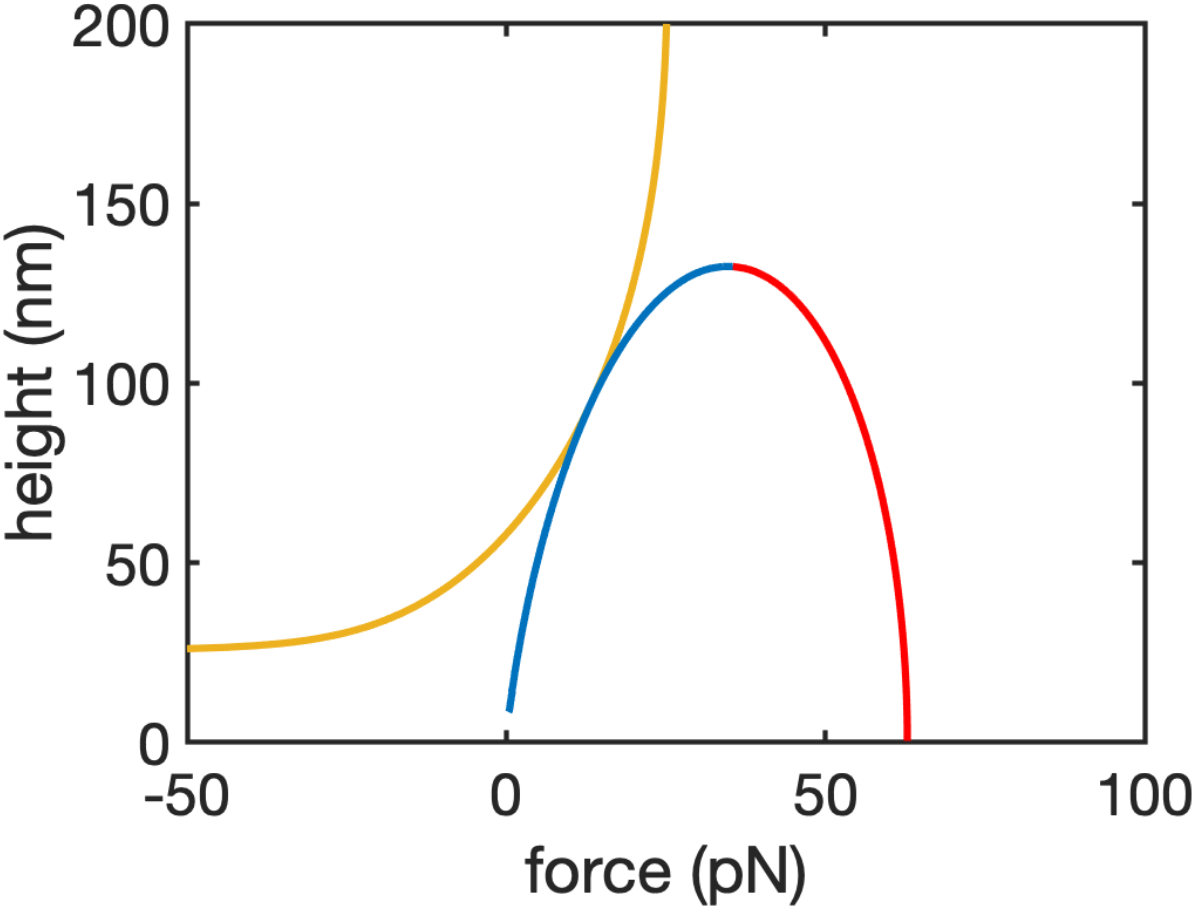
Force vs height for the fusion pore families with freely hinged boundary conditions. Membrane tension is 0.1 pN/nm and the membrane diameter *D*_mb_ is 200 nm. Catenoids (red and blue) require a pulling (positive) force to maintain the height, while for tethers of small heights compressive forces are required.

**Figure S2.**
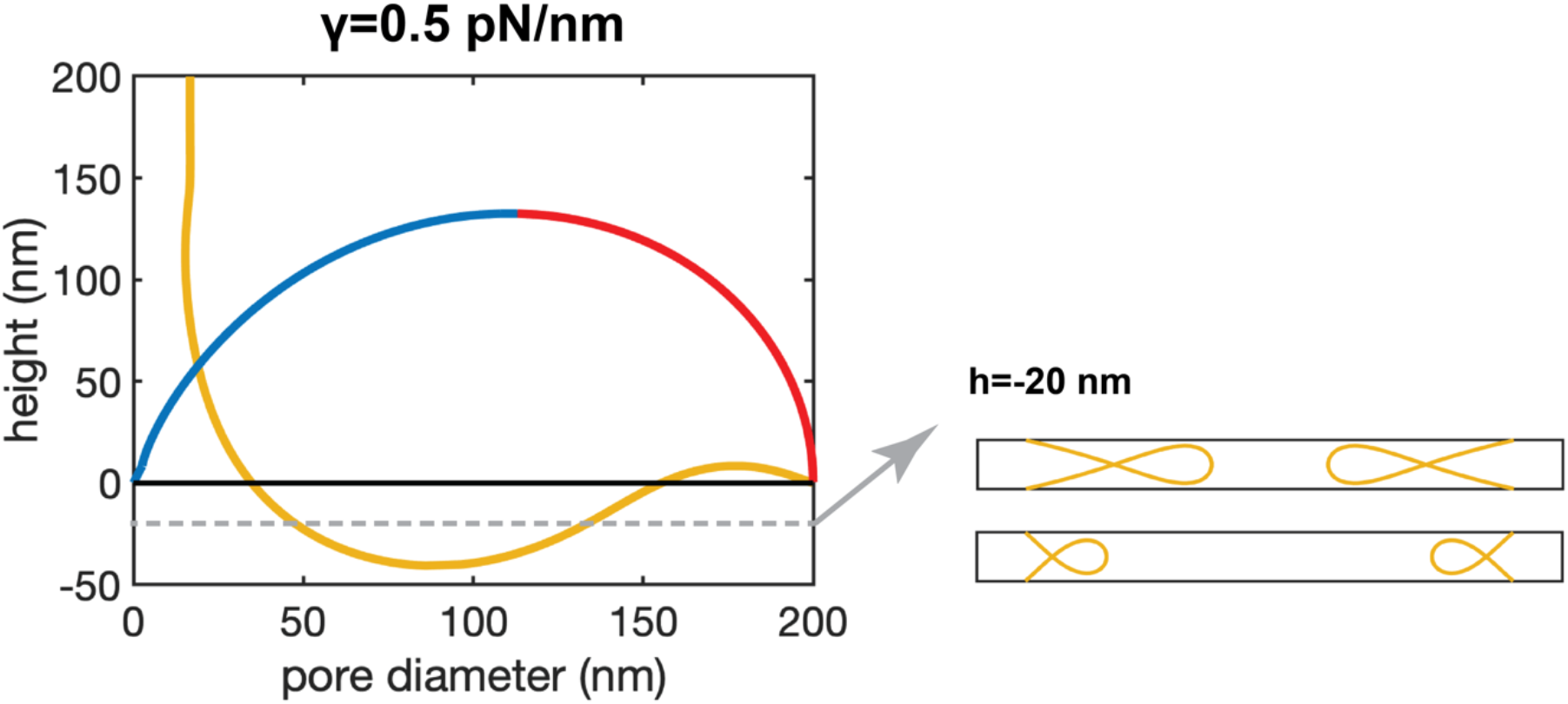
Large tether oscillations under a higher membrane tension. Under freely hinged boundary conditions, the tether family oscillates to negative separations. However, these solutions are not physical.

**Figure S3.**
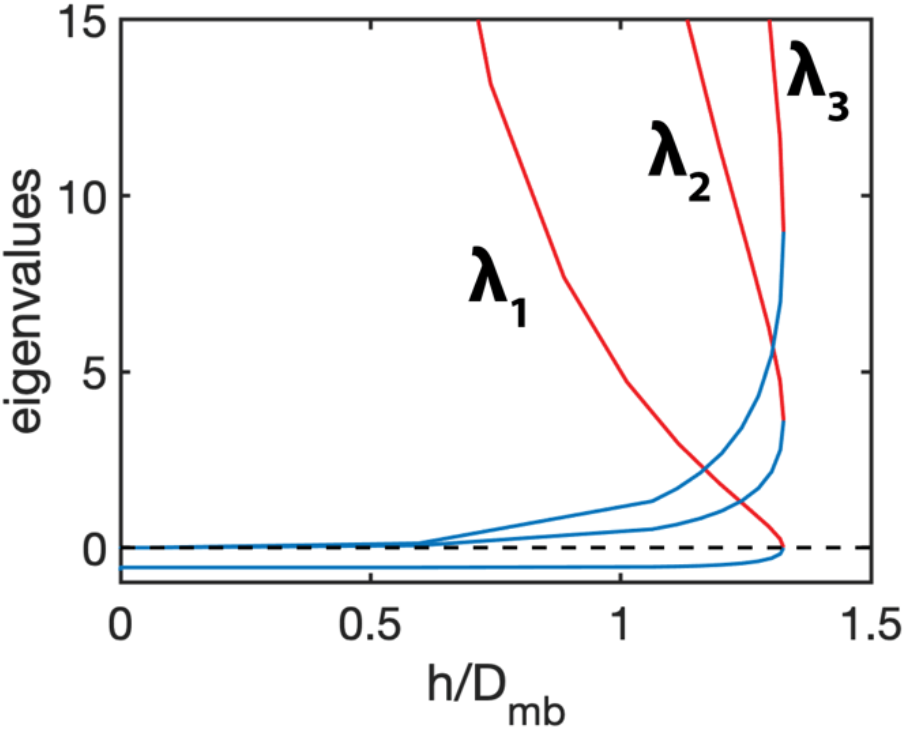
The smallest three Eigenvalues of Equation 8. For thin catenoids (blue), the first (smallest) eigenvalue *λ*_1_ is negative but other eigenvalues *λ*_2_, *λ*_3_,… are always positive. For wide catenoids (red), all the eigenvalues are positive.

**Figure S4.**
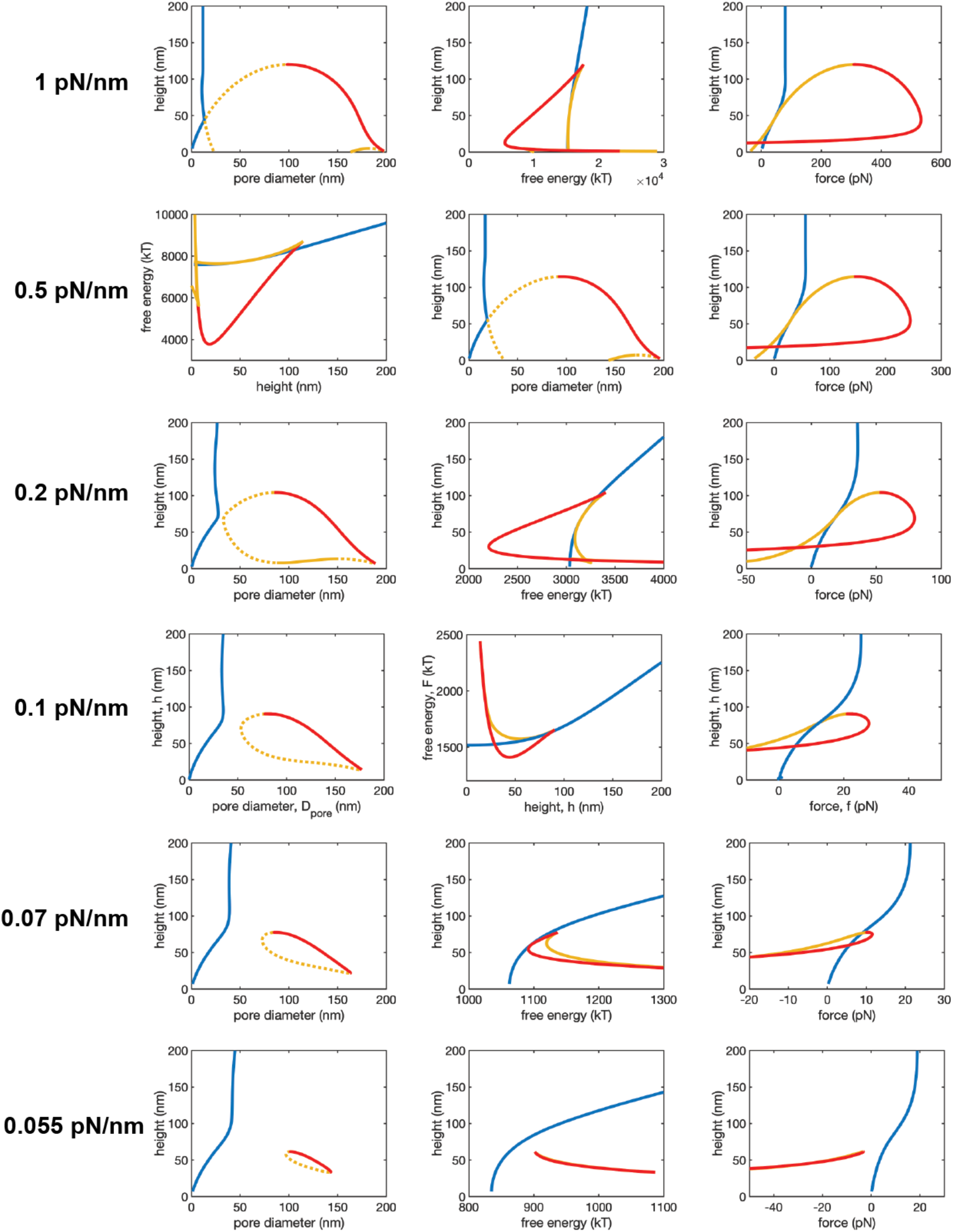
Pore diameter, free energy and force of the three fusion pore families vs pore height under clamped boundary conditions. Membrane diameter is 200 nm. With decreased membrane tension, the wide quasi-catenoids (blue) only exist in a smaller range of heights, and below a certain tension, the thin quasi-catenoids (red) starts to have the lowest energy.

**Figure S5.**
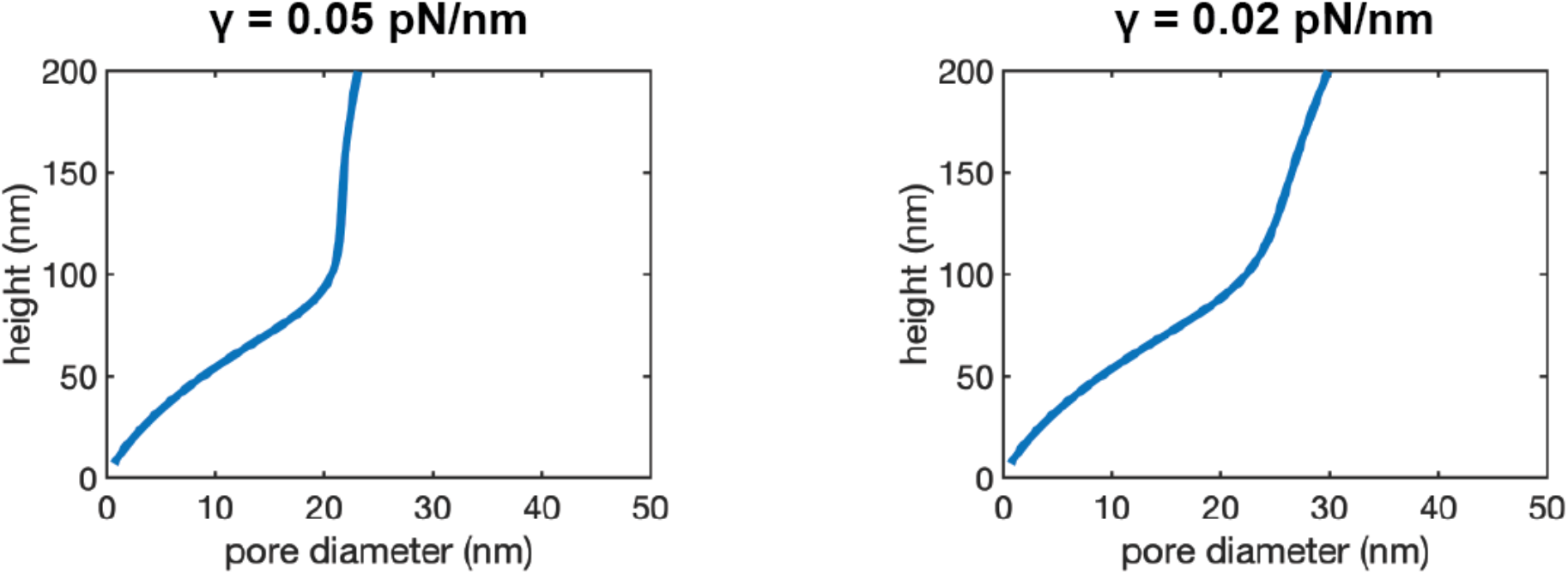
Thin quasi-catenoid is the only realizable pore under low membrane tension and clamped boundary conditions. With membrane diameter is 200 nm, only the thin catenoid pores are found, consistent with Fig. S4 where the other two families, wide catenoid and the unstable intermediate, exist in a smaller and smaller height range with decreased tension.

**Figure S6.**
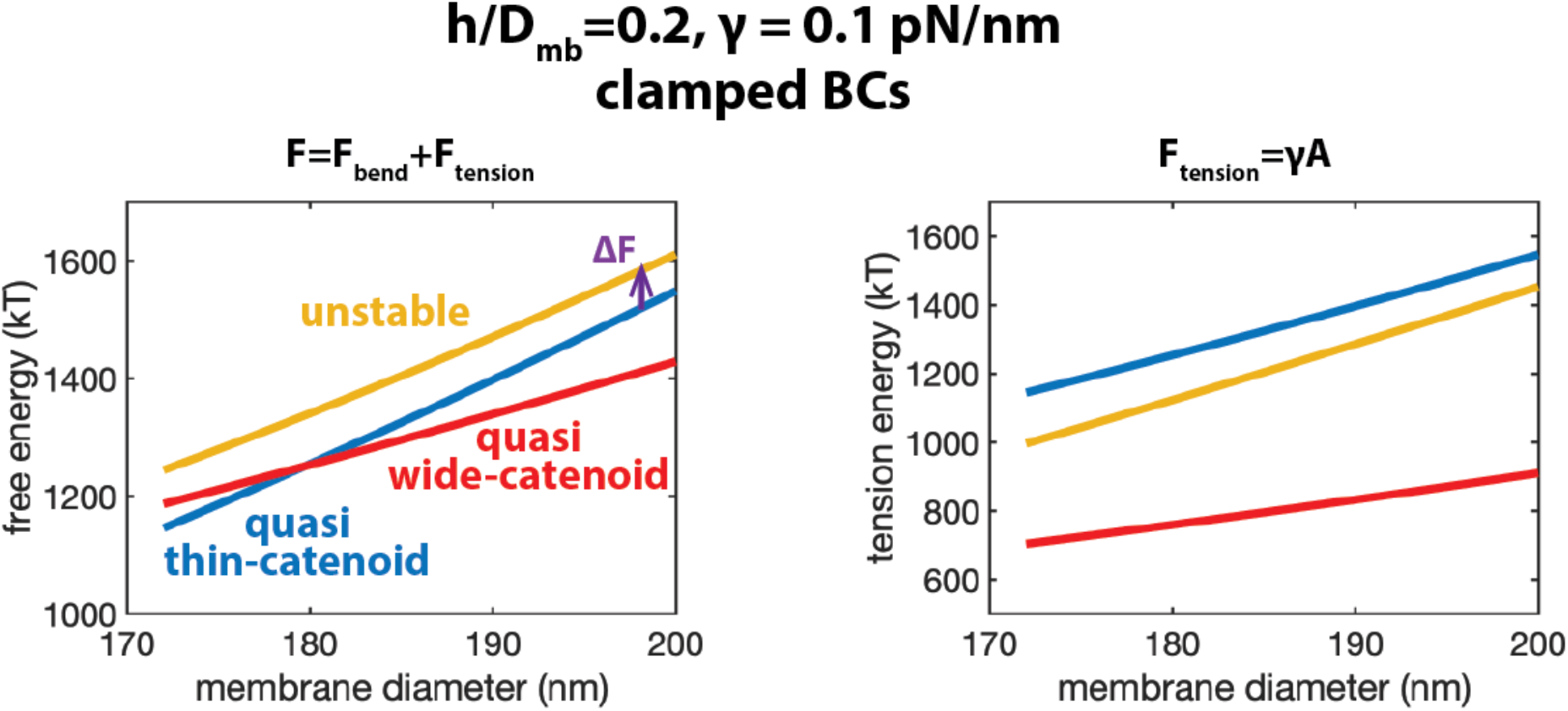
Pore dilation is favored by larger vesicles. The free energies of the three families of fusion pore all increase with increased membrane diameter, but at different rate. The energy of the thin quasi-catenoid increases the most, mainly attributed to the large area increase (right). The energy of the unstable intermediate only increases moderately, thus the energy barrier for pore dilation Δ*F* is smaller with increased vesicle size.

**Figure S7.**
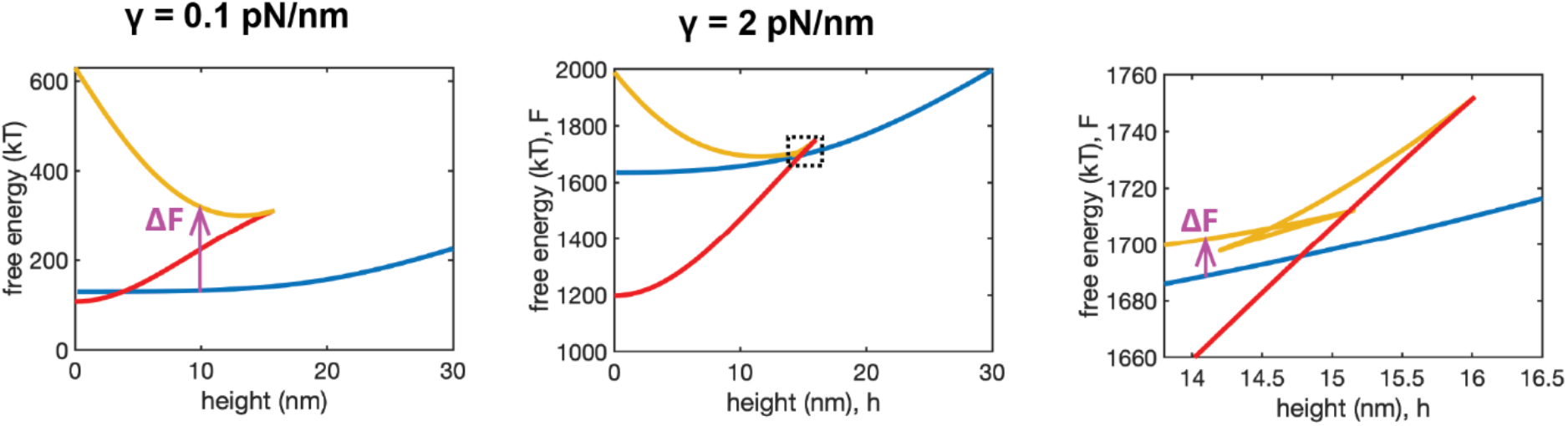
Nanodiscs lock the fusion pore into the thin quasi-catenoid family but high tension may activate pore dilation. Under normal tension (left), the transition from the thin catenoidal pore (blue) to the unphysical wide catenoidal pore (red) is blocked by a high energy barrier ΔF of ~150 kT. However, under high tension (middle), the barrier decreases to ~10 kT (right, zoom-in of the dotted box in the middle panel) which makes the unphysical wide catenoidal pore realizable.

